# Combination of favorable gene haplotypes plays an important role in rice hybrid grain quality traits based on genome-wide association analysis

**DOI:** 10.1101/2020.06.04.134023

**Authors:** Lanzhi Li, Xingfei Zheng, Xueli Zhang, Kai Xu, Shufeng Song, Jing Su, Chao Wu, Zirong Li, Wenbang Tang, Ying Diao, Zhonghai Tang, Zheming Yuan, Zhongli Hu

## Abstract

Yield level and grain quality determine the commercial potential of rice (*Oryza sativa* L.) varieties. Mining and using genes that control important rice grain quality characteristics are major tasks for plant breeders. Here, a genome-wide association study was conducted to determine the genetic bases of 12 rice grain quality traits in 113 varieties and 565 testcross hybrids. A total of 56 significant SNPs were associated with 9 of the traits in variety phenotypic, general combining ability, testcross hybrid phenotypic and mid-parental heterotic datasets, from which 45 novel loci were identified. The cumulative effects of superior alleles or favorable haplotypes of genes closest to significant quality trait-associated loci were found in the four datasets. Additionally, the favorable gene haplotypes performed better than those of superior alleles in the variety and testcross hybrid datasets. Pyramiding the favorable haplotypes of five cloned rice grain quality genes resulted in a very low amylose content and high yield in the latter. These testcross hybrids had rice grain qualities similar to their parental lines but with much higher yields. The amylose content, grain width and grain length predictions for potential hybrids among the 113 varieties using genomic selection based on the BayesB method revealed a performance trend similar to those the testcross hybrids in our study. Thus, the selection of combination of favorable rice quality-related gene haplotypes is recommended to breed hybrids with high yields and elite grain qualities.

## Introduction

More than half of the world’s population consume rice as their staple food. With improvements in living standards, people are paying more attention to food quality and a healthy lifestyle, which requires rice (*Oryza sativa* L.) to have a higher end-use cooking quality (Huggins et al., 2018). Grain quality is a complex trait regulated by interacting genetic and environmental factors. Mining and utilizing genes that control important quality characteristics of rice are major tasks of plant breeders (Huang et al., 2010; McCouch et al., 2010; Tung et al., 2010). However, the simultaneous improvement of yield and end-use quality remains a challenge for rice breeders because yield and quality are typically negatively correlated (Huggins et al., 2018). In general, the yields of hybrid rice are greater than those of conventional rice varieties, but the grain quality is not improved. To obtain both high-quality and high-yield rice varieties, the quality traits of hybrid rice need to be genetically dissected.

A high milled brown rice rate, translucent grains and good taste are desired on essentially all markets, and these grain characteristics are the bases of evaluation indexes for cooking rice. Rice grain quality has been classified into milling, appearance, cooking and nutritional categories (Fitzgerald and Resurreccion, 2009; Huggins et al., 2018). In recent years, with the rapid development of molecular biology-based theories and technology, many rice grain quality trait-related genes have been cloned and localized, and their mechanisms and functions have also been investigated. Having genomic markers linked with various grain quality traits in rice will help breeders combine crop productivity with desirable end-use quality traits targeted for specific markets (Huggins et al., 2018). With molecular marker-assisted selection, breeders can introduce good quality genes into high yield varieties and greatly improve the breeding efficiency.

Grain size is closely related to the yield (Che et al., 2016; Asante, 2017). At present, dozens of grain size-related genes have been isolated from different rice germplasm resources, such as the genes *GS3, GL3.1* and *GW7/GL7*, controlling grain length (GL; Mao et al., 2010; Wang et al., 2015a; Wang et al., 2015b), the genes *GW2, GW5/qSW5* and *GS5*, controlling grain width (GW; Song et al., 2007; Wang et al., 2008; Li et al., 2011), and the genes *GS6, GS9, TGW6* and *GW8/SPL16*, controlling grain size (Sun et al., 2013; Ishimaru et al., 2013; Song et al., 2015; Che et al., 2016). Chalky grains are considered as low quality because of their poor appearance and the undesirable cooking and milling qualities (Zhao et al., 2018). The rice gene *OsRab5a* regulates endomembrane organization and storage protein trafficking in rice endosperm cells, which affects the formation of amyloplasts (Wang et al., 2012); and *Chalk5* encodes a vacuolar H^+^-translocating pyrophosphatase, which influences grain chalkiness in rice (Li et al., 2014).

The amylose content (AC) has the greatest influence on the cooking and consumption qualities of rice. The synthesis of rice amylose is catalyzed by granule-bound starch synthase protein that is encoded by the waxy gene *Wx* (Smith et al., 1997). There are several alleles of the waxy gene, including *Wx, Wx*^*a*^, *Wx*^*b*^, *Wx*^*mp*^, *Wx*^*op*^, *Wx*^*in*^ and *Wx*^mq^. Zhou et al. (2016) cloned a practical resistant starch gene based on the physical map. Rice varieties with an intermediate gel temperature, which is predominantly determined by amylopectin structure, are generally preferred by consumers. The gene, starch synthase II is the major determinant of gel temperature (Asante, 2017).

Protein content (PC) is a major index of rice grain nutritional quality. Generally, the PCs in *indica* rice varieties are higher than those in *japonica*. In total, 19 QTLs have been linked to rice grain PC. Liao et al. (2019) found that over-expressed *OsAPP6* could facilitate the roots absorption of NH4^+^, which improved rice grain protein synthesis.

QTL mapping and cloning provide important information for dissecting main-effect genetic loci, but it is often difficult to mine the minor-effect genes associated with traits. In addition, only the genetic information of the two parents are included in a QTL analysis, making it difficult to identify the network relationships among complex genes in narrow germplasm resources (Huang et al., 2018, Fang et al., 2018). A genome-wide association study (GWAS) has a higher resolution than QTL mapping because it includes the recombination information accumulated by natural populations during long-term evolution. Fine positioning of the QTL can be achieved in an association analysis, even directly targeting the gene itself (Huang et al., 2018). GWASs have been widely applied in crops, such as wheat, corn, potato, adzuki bean, perennial ryegrass, rice and mustard, to decipher the genetic bases of quantitative traits (Huang et al., 2010; Xie et al., 2010; Huang et al., 2015; Huang et al., 2016; Huang et al., 2018),

Zhao et al. (2010) used a chip containing more than 36,000 high-quality SNPs to perform a GWAS on 34 traits in more than 400 rice varieties in 82 countries. Some known genes were found, such as *qSW5* and *GS3*, which control grain size. With the development of high-throughput sequencing technology, GWASs based on whole-genome sequencing have been widely used in research on rice agronomic and quality traits. In 2015, Huang et al. directly sequenced 1,495 hybrid rice varieties in China’s main rice producing areas and used 38 agronomic traits, such as yield, quality and disease resistance, of these hybrid rice varieties. In 2018, Misra et al. conducted a GWAS on cooked rice’s quality traits (adhesion, hardness, elasticity, viscosity and AC) for 236 *indica* rice varieties from 37 countries. A total of 224 significantly associated SNP loci were detected, of which 97 were validated by at least two GWAS methods. GWAS of three rice diversity panels were performed by Huggins et al. in 2018. Their results revealed large- and small-effect loci associated with known genes and previously uncharacterized genomic regions and suggested that multiple grain quality traits were inherited together.

At present, GWAS research on rice quality traits is mainly conducted using conventional rice accessions, while the study of hybrid rice quality is limited. To obtain both high-quality and high-yield rice varieties, it is necessary to decipher the genetic bases of hybrid rice grain quality. In this study, a GWAS analysis was performed on the grain quality traits of 113 *indica* rice varieties in China and their 565 testcross hybrids based on the North Carolina II (NCII) design. The aim of this study is to provide greater insights into effective breeding strategies for the grain quality improvement of rice hybrids.

## Results

### Rice grain quality trait phenotypes and their relationships

Phenotypic distributions of the 12 quality traits in testcross hybrids and their male parental lines are shown in Fig. 1a and Supplementary Figure 1 and 2. The 12 traits in the four datasets (V, TC, Gca and Hmp) were distributed continuously with diverse variation levels, indicating that complex genetic mechanisms underlie rice grain quality traits. Rice grains with low CD and PGWC, and high TD, PC and ASV, are desired for elite quality rice. In this study, the mean values of CD and PGWC in the testcross hybrids were higher than those of their male parental lines, but the former had lower TDs, PCs and ASVs. This result partially confirmed that hybrids have poor rice grain qualities when compared with those of conventional varieties. For the remaining seven traits, testcross hybrids showed similar performances with those of their male parental parents.

**Figure 1.**
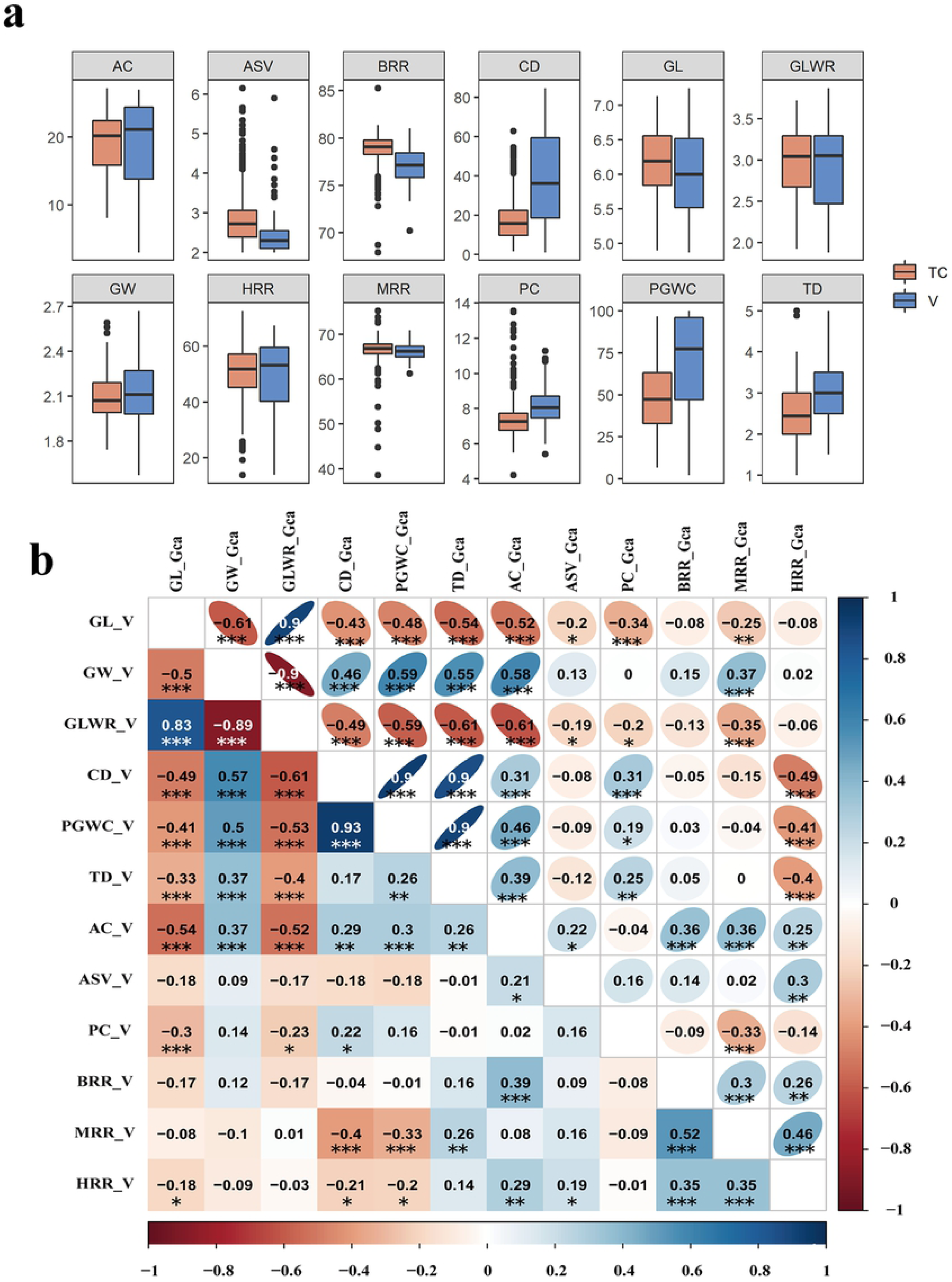
Phenotype distributions and phenotypic correlations among different traits. (a) Phenotype distributions of rice grain quality traits in Varieties (V) and TC (Testcross) dataset. (b) Phenotypic correlations among different traits. The left bottom panel represents the phenotypic correlations in the V dataset, and the right top panel represents the phenotypic correlations in the Gca dataset. The values indicate correlation coefficients (*r*). The areas and colors of ellipses represent the absolute values of the corresponding *r*. Right and left oblique ellipses indicated positive and negative correlations, respectively. Asterisks indicate significant correlations using a two tailed *t*-test (\**p* < 0.05, \*\**p* < 0.01 and \*\*\**p* < 0.001).

Correlation analyses for every pair of traits were performed within the four datasets. The trends of most traits’ relationships were consistent within the V and Gca datasets (Fig. 1b), and within the TC and Hmp datasets (Supplementary Figure 3). Additionally the significance levels of the correlations varied among traits in the same dataset. GL and GW are closely related to yield, and these two traits are always negative correlated. In this study, GL was significantly negatively correlated with CD, AC and TD in all the datasets, while GW was positive correlated with them. Generally, grains with a low CD, long GL and low AC are desired in rice. The correlation coefficient analysis results confirmed the trend and indicated that the germplasm materials in this study may have been preserved by long artificial selection. We found that GW and GL were not significantly associated with ASV, BRR, MRR or HRR. Thus, breeders should consider the relationships among traits when conducting rice breeding for high yield and grain quality.

### Structure of the NCII population

The NJ tree suggested that there were population differences within the male parental lines (Supplementary Figure 4); however, no clear clades were shown, which indicated that these materials have a complicated breeding history. The male sterile lines and restorer lines were not separated into two divergent clusters, which further indicated the wide variation within our materials. Five branches of testcross hybrids were detected at *K* = 5 (Supplementary Figure 5). The testcross hybrids derived from the same male parent line clustered in the same clades in the NJ tree, which was similar to the findings of Huang et al. (2015) and Zhen et al. (2017). Therefore, when performing a GWAS analysis of TC and Hmp, special attention should be paid to the influence of population structure. The PCA of the 120 parental lines revealed a wide variation among them (Supplementary Figure 6), while a PCA revealed a highly-structured genetic relationships among the TC lines (Supplementary Figure 7). The first and second genomic PCs mainly separated the TC lines into two groups, while the third and fourth PCs separated them into four small groups consistent with their female parental origins, except for 3A and Y58S, which were together. To correct for subpopulation structure, we integrated the first three PCs as covariates within the model and performed a GWAS across all the accessions in the V, Gca, TC and Hmp datasets.

### Whole-genome screening for significantly associated loci

To reduce false positive or negative results, four algorithms: general linear model, mixed linear model, compressed mixed linear model-based EMMAx (Kang et al., 2010) and fixed and random model circulating probability unification model in software GAPIT were performed in the GWAS pipeline. Then, the Q-Q (quantile-quantile) plot and the Manhattan graphs obtained by each model (Figs 2 and 3, Supplementary Figures. 8–31) were compared to determine the best results for each trait in all the datasets (12 × 4 = 48 datasets in total), which were used for subsequent analyses (Supplementary Table2). Using a strict Bonferroni correction, the threshold was set to *p* < 2.64 × 10^−8^ (0.01/total number of SNPs), and 56 significant SNPs were detected, involving 9 traits and 24 datasets (Table 1).

**Table 1.**
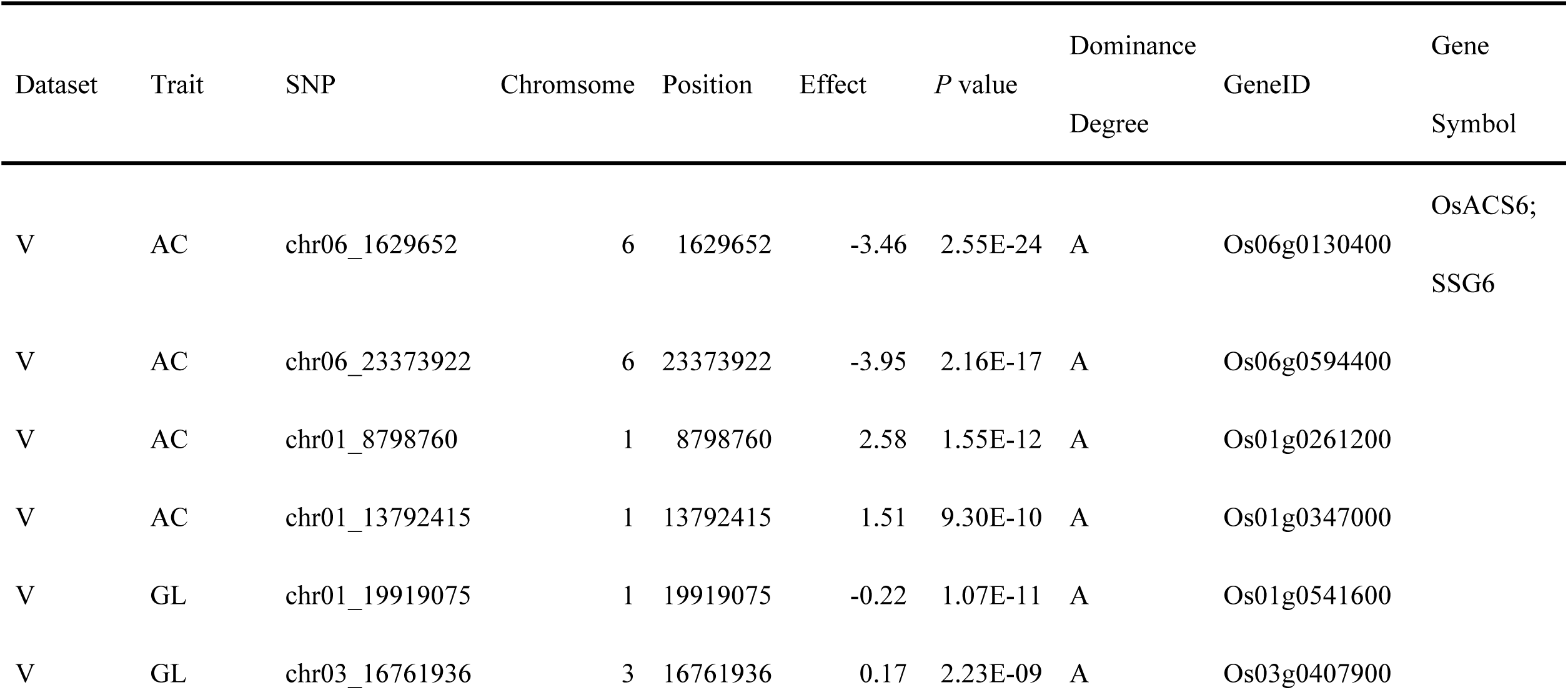
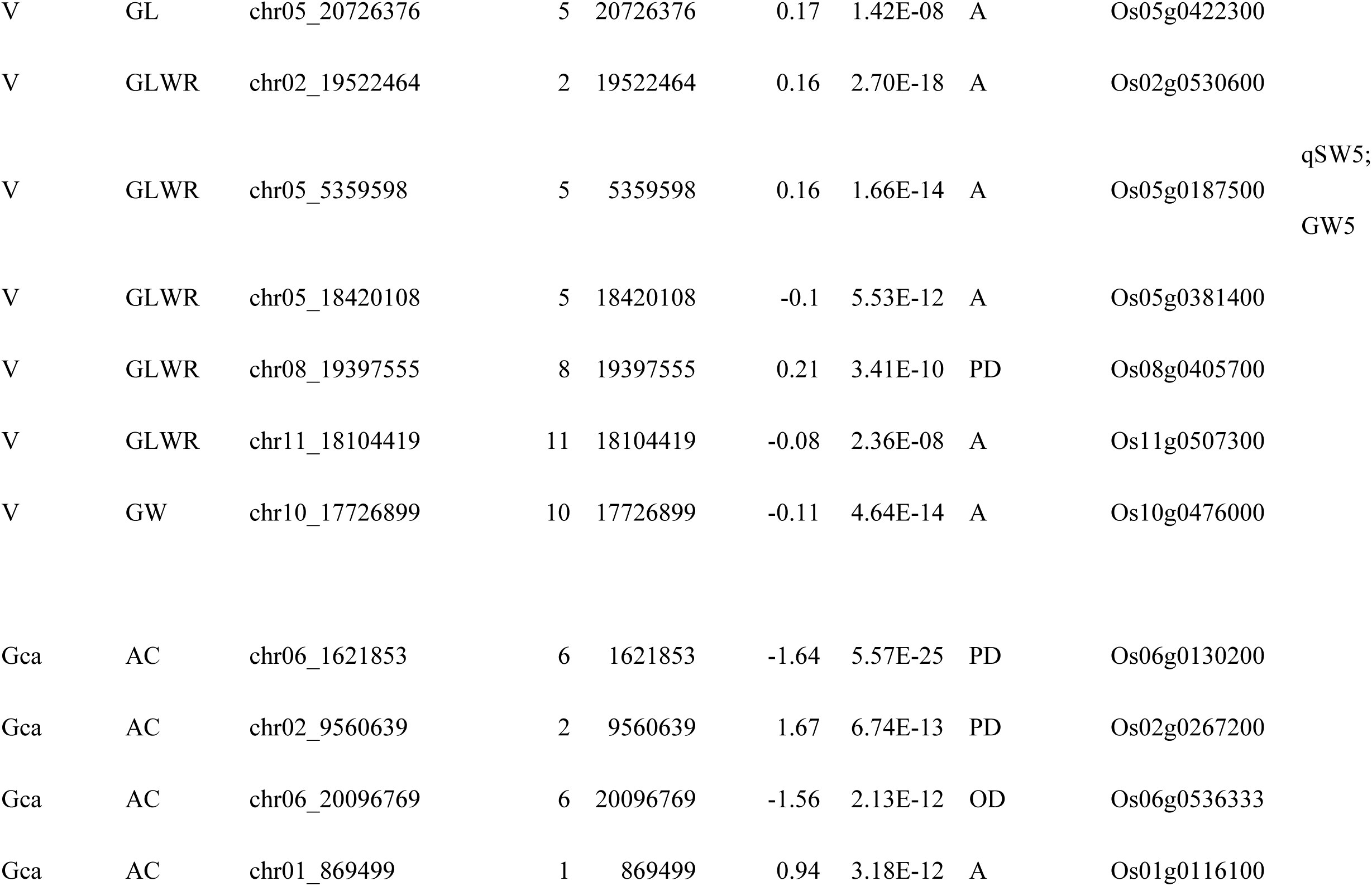
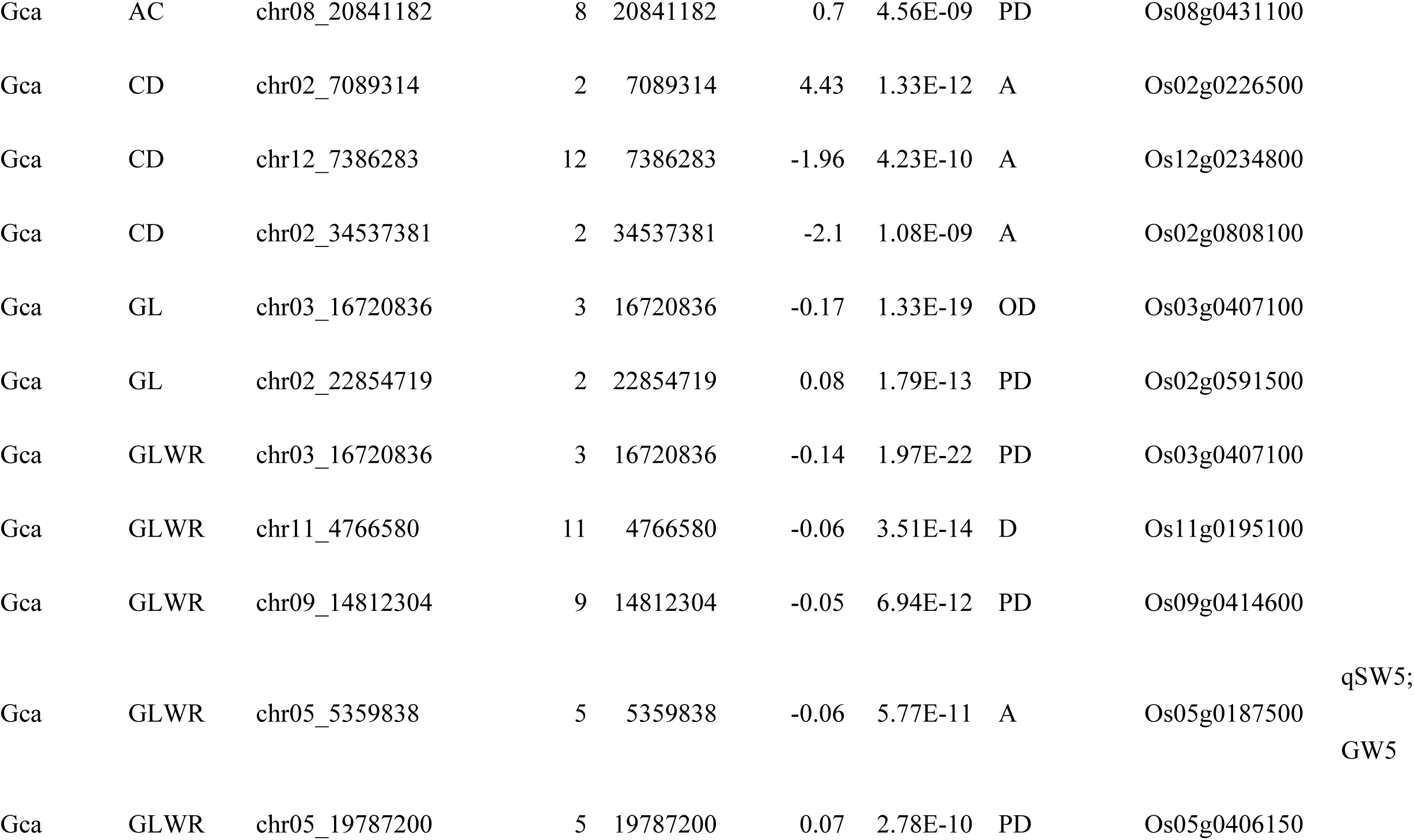
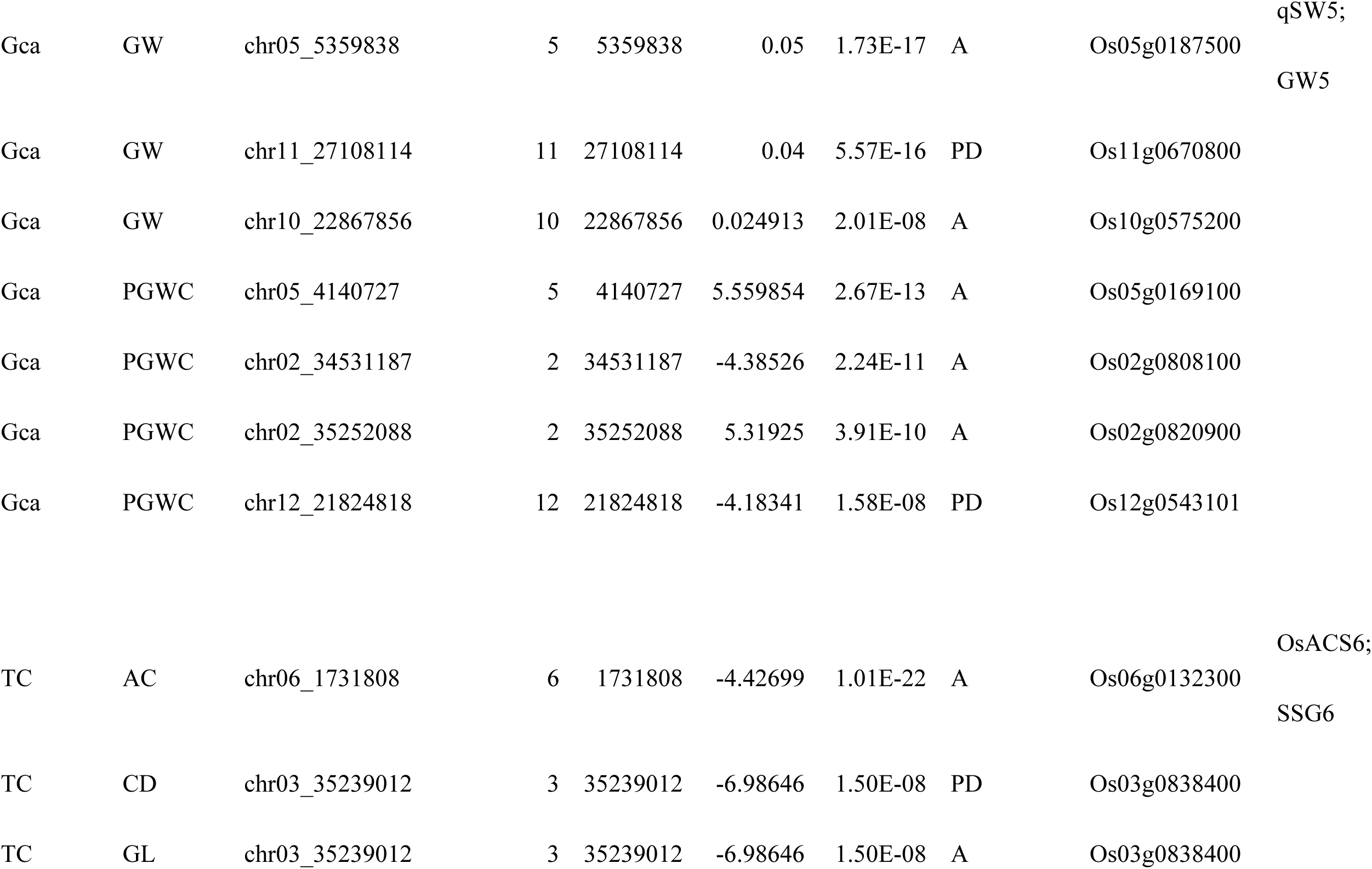
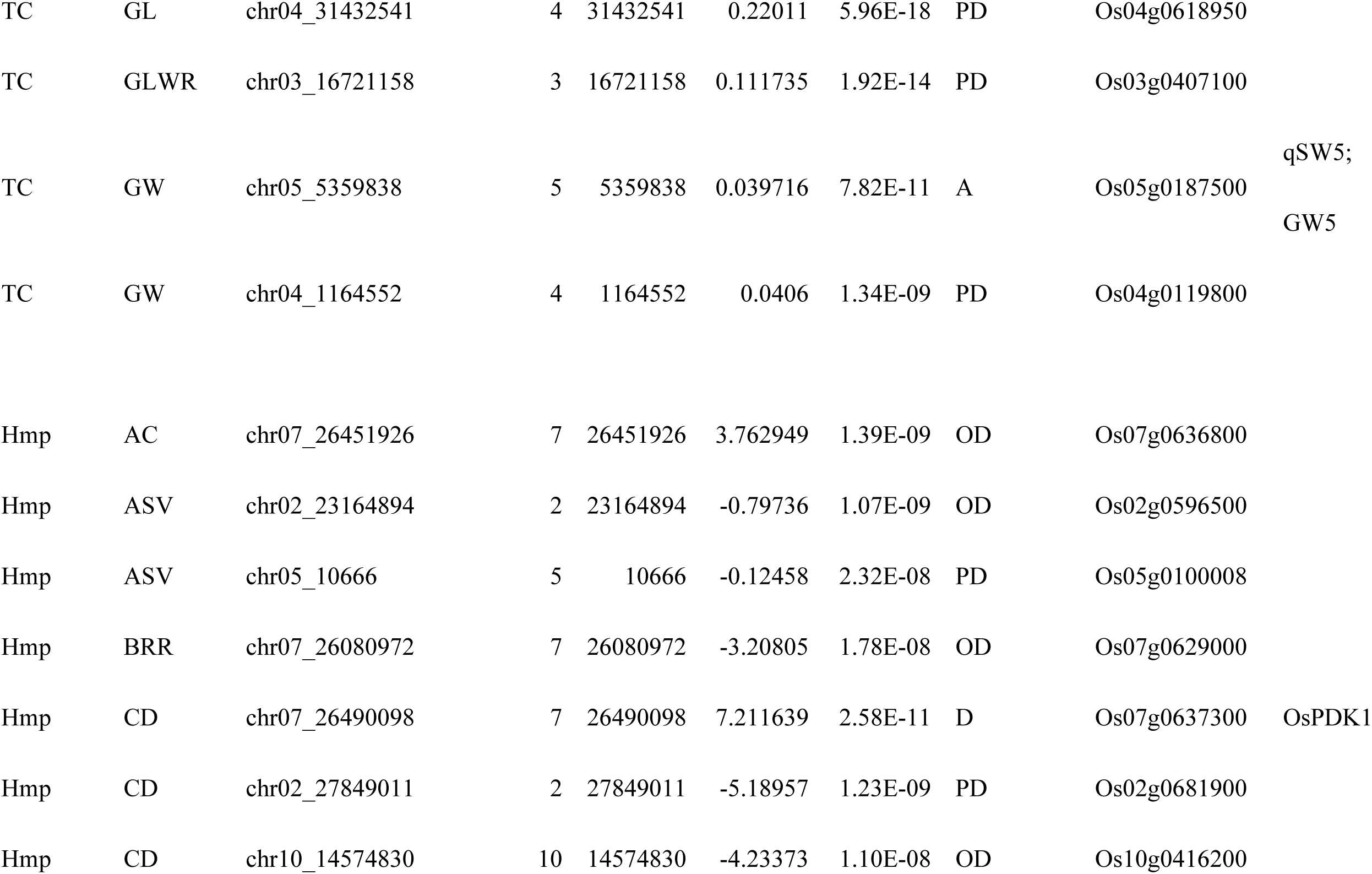
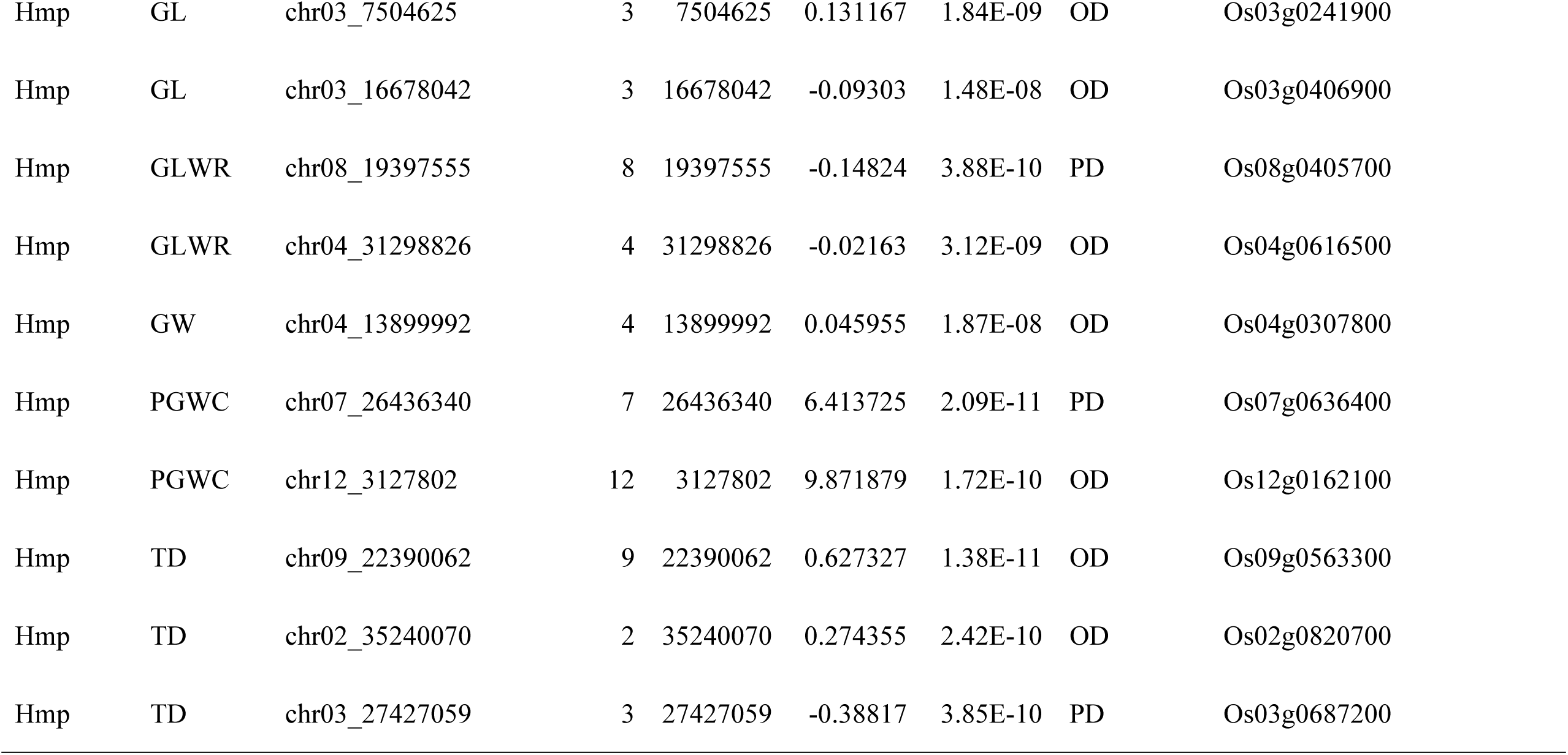
Significant markers associated with rice grain quality traits in four datasets.

**Figure 2.**
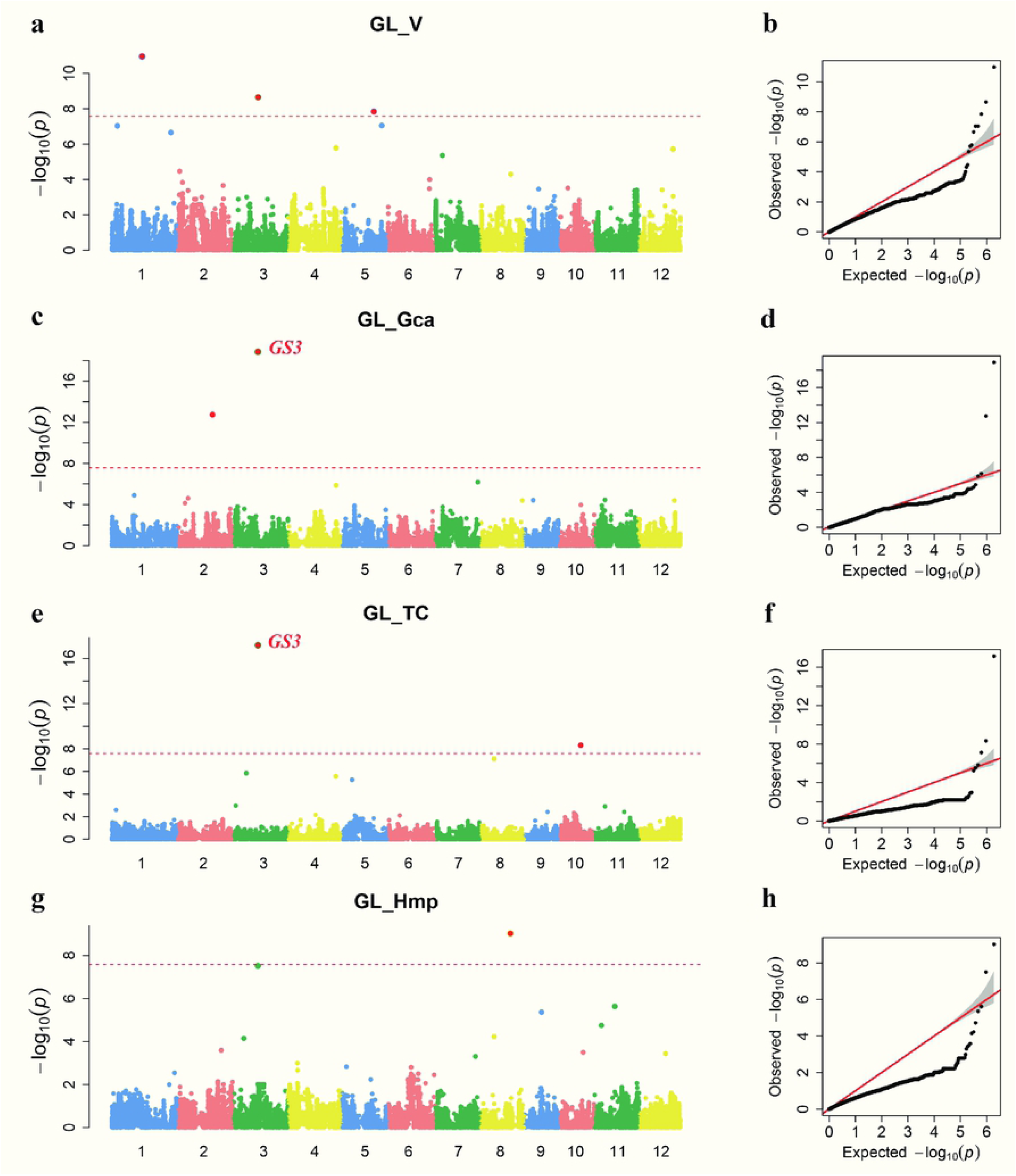
GWAS results for GL. (a, b) Manhattan plots and and quantile–quantile about V of GL. (c, d) Manhattan plots and and quantile–quantile about Gca of GL. (e, f) Manhattan plots and and quantile–quantile about TC of GL. (a, b) Manhattan plots and and quantile–quantile about Hmp of GL. The genome-wide significant P value threshold *P*<2.64*10^−8^ is indicated by a horizontal dash-dot line. Significant SNPs with *P*<2.64*10^−8^ are depicted as red dots. The well-characterized genes near significant loci are indicated in red characters.

**Figure 3.**
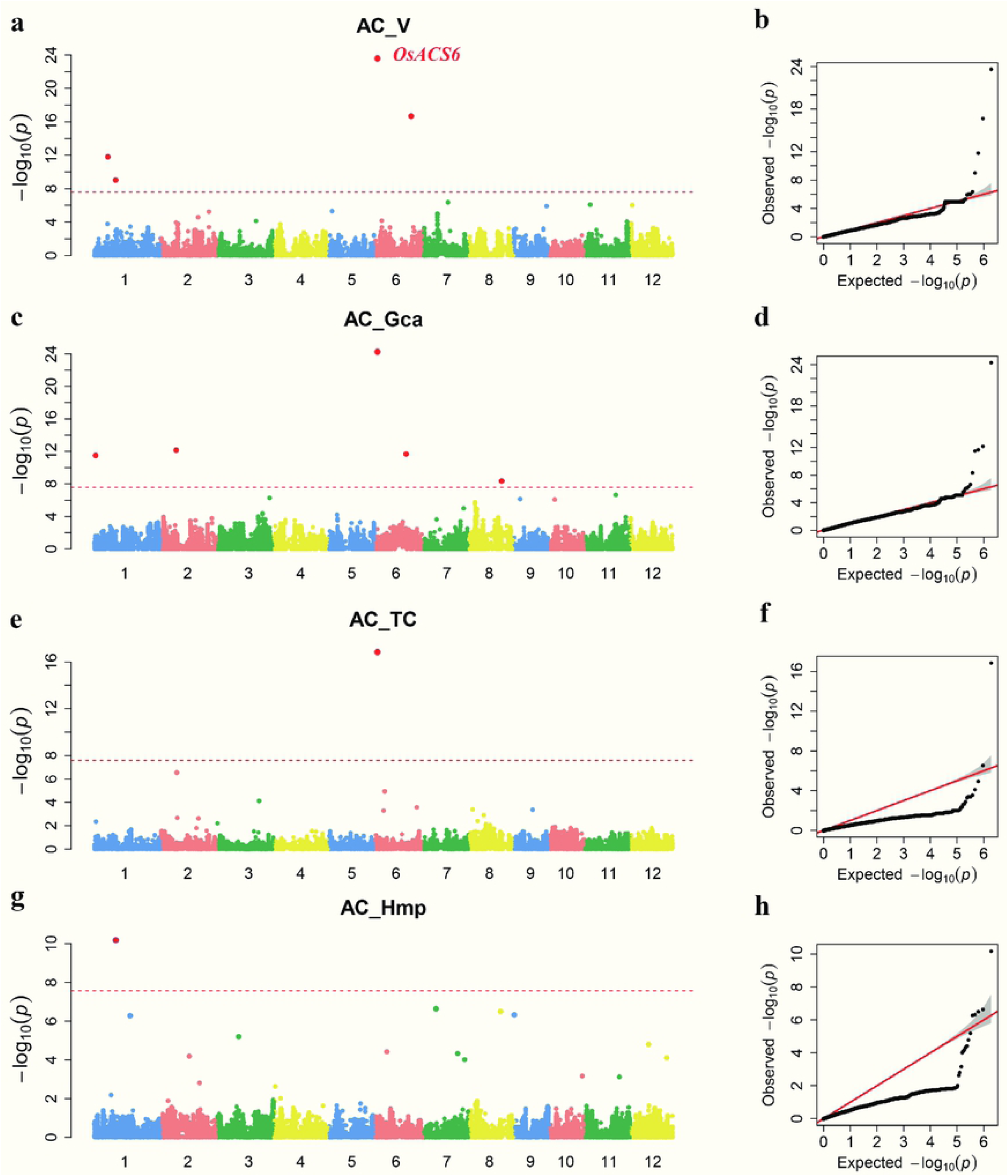
GWAS results for AC. (a, b) Manhattan plots and and quantile–quantile about V of AC. (c, d) Manhattan plots and and quantile–quantile about Gca of AC. (e, f) Manhattan plots and and quantile–quantile about TC of AC. (a, b) Manhattan plots and and quantile–quantile about Hmp of AC. The genome-wide significant P value threshold P<2.64*10^−8^ is indicated by a horizontal dash-dot line. Significant SNPs with P<2.64*10^−8^ are depicted as red dots. The well-characterized genes near significant loci are indicated in red characters.

In our research, four of eight significant markers detected for GL harbored or were located near reported genes (*GS3, GL3.2, GL7, PGL1* and *PGL2*) associated with GL (Fig. 2). For GW, SNP chr05_5359838 was simultaneously detected as being significant in Gca and TC. It was located near the GW main-effect gene *GW5*. SNP chr03_35360872 was harbored in gene *qTGW3*, which regulates rice GW (Table 1). The waxy gene is involved in amylose synthesis in the endosperm and is located on chromosome 6. Approximately 34 kb apart from it, the significant SNP chr06_1731808 was identified for AC in the TC dataset (Fig. 2). For chalkiness, SNP chr05_4591773 for CD was detected as being significant in the Gca and TC datasets. It was approximately 120 kb from the *chalk* gene, which is the main-effect gene for grain chalkiness regulation. SNP chr05_4140727 for PGWC in the Gca dataset was 80 kb from *chalk*. The other 49 significant markers for grain quality traits were first detected here.

### Assessing pleiotropy between traits

Pleiotropy and LD play important roles in identifying correlations among phenotypes. In this study, we found that several SNPs or their nearby regions associated with different traits (Supplementary Table 3), which helped us dissect the genetic architecture of the correlations across different traits. For example, the chr03_16720836 locus was simultaneously detected to be correlated with GL in Gca, GL in TC and GLWR in Gca, and 0.32 kb from this locus, SNP chr03_16721158 was detected to be correlated with GLWR in TC. These two loci are in the vicinity of the cloned gene *GS3* (position: chr03:16729501 to 16735109), which is the major gene controlling rice GL and GW, and also a minor-effect QTL controlling rice GW and grain filling. In addition, the same traits in different datasets had overlapping peaks within a given genomic region. SNP chr06_1621853 on chromosome 6 associated with AC in Gca, which is located at a close distance from SNP chr06_1629652, which was significantly associated with AC in the V dataset. *OsACS6*, which regulates the size and morphology of starch granules in endosperm cells, is located near these two sites. These results are not only consistent with other researcher’s findings, but also suggest that some traits share a genetic basis that allows a general combing ability or heterosis. Some quality traits, or the same trait’s general combing ability and heterosis, are more likely to be co-inherited, and these important loci may be used in breeding to improving hybrid rice quality.

### Dominance degree

To dissect the genetic basis of rice grain quality trait heterosis, we calculated the dominance degree of the top 100 SNPs of the GWAS results for all the data using descending −log (*P* values). Generally, the dominance degree varied in the four datasets. Except for PGWC and PC, all the traits showed A followed by PD in the V dataset (Supplementary Figure 32). In the Gca dataset, on the whole, A and PD accounted a greater proportion than D and OD. A accounted for a large proportion of the traits GW, ASV, MRR and HRR, while PD mainly accounted for GL, PGWC, CD and BRR. Similar trends in the dominance degrees of quality traits occurred in the Gca dataset. The four types of dominance existed and varied among traits, but in general, A and PD were still relatively large in the TC dataset. The OD proportion increased in the Hmp dataset when compared with TC dataset. Compared with the phenotypic values of the testcross hybrids, the Hmp value is more suitable for detecting the heterozygotic genetic basis. Because only dominant effect involved in Hmp at single locus level, while additive and dominant effect in testcross hybrid phenotypic performance (Li et al., 2008). Thus, A is main genetic basis of rice quality traits in these accessions (male parental lines in this study), while OD mainly underlies the heterotic genetic basis.

### Frequency of superior alleles of associated loci and favorable haplotype of gene

The frequency of superior alleles of associated loci in V and TC datasets was investigated following the method of Huang et al. (2015), in which the allele with the better quality performance (for example, greater GL, greater GW or lower AC) was defined as the superior allele. Because of the low LD rate of decay in rice, a GWAS cannot resolve a single gene (Huang et al., 2010), we investigated the frequency of the favorable haplotype of the closest gene to the associated loci. Similarly, the haplotype having the better quality performance was defined as the favorable haplotype of the gene. Our results (Fig. 4) showed that, overall, the performance of paternal lines or testcross hybrids of each trait improved as the number of superior alleles or favorable haplotypes increased. Additionally, the performance of the favorable haplotype of the gene outperformed the superior allele of the SNP (lower AC, similar GW and longer GL) in the V dataset. The TC lines in which favorable haplotype of genes were pyramided also showed good rice grain quality that approached those of the male parental lines but much higher yields, which suggested that breeders may select hybrids having high yields and elite quality.

**Figure 4.**
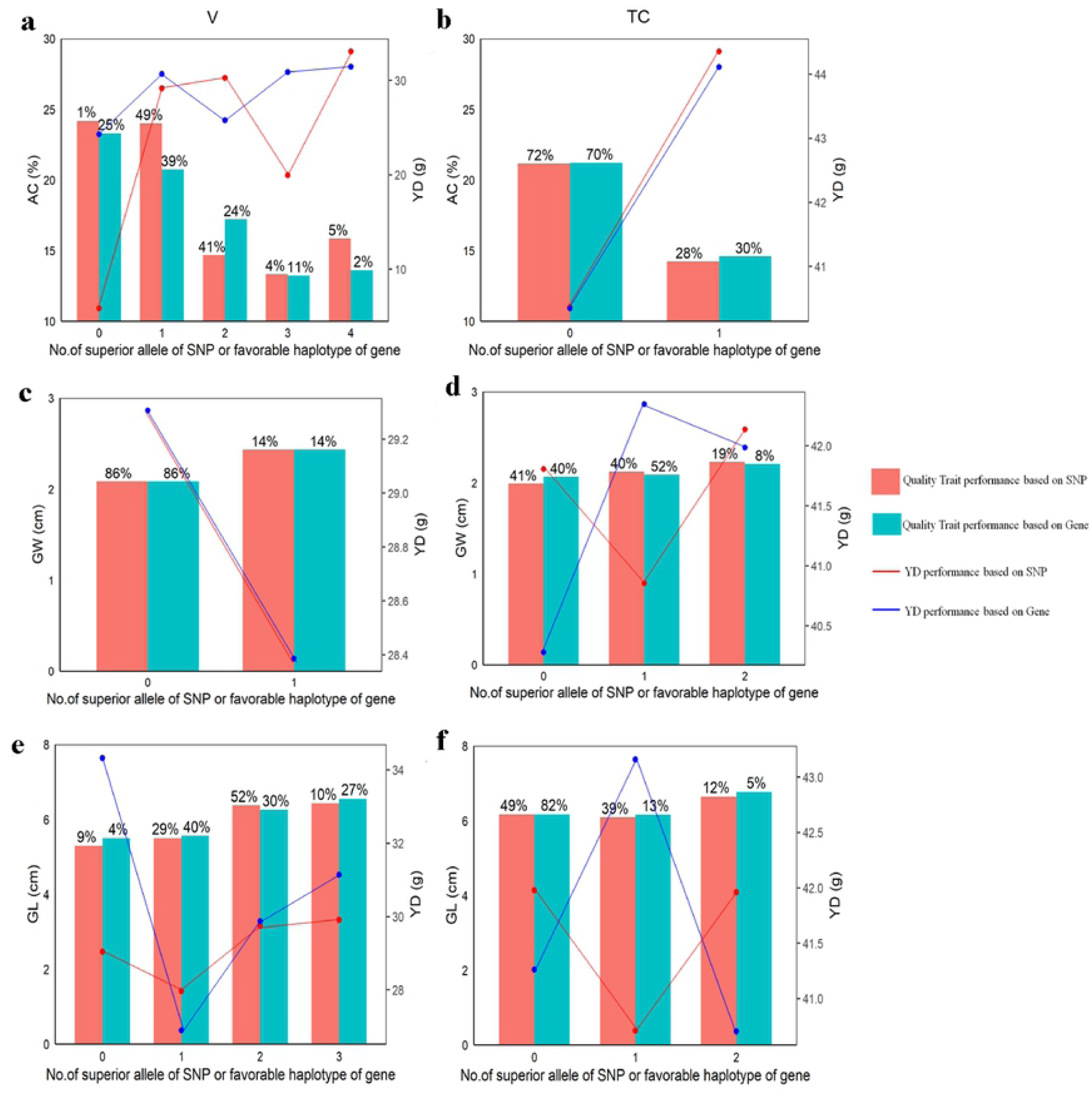
The phenotype values and the percentage of accessions with different number of superior alleles of significant loci or favorable haplotype of closest gene of significant loci for quality trait in V and TC datasets. a, c, e separately represents AC, GW and GL with different number of superior alleles or favorable haplotype of genes in V dataset, b, d, f separately represents AC, GW and GL with different number of superior alleles or favorable haplotype of genes in TC dataset. Percentage showed on bars are rate of corresponding lines to whole lines in V or TC dataset.

### Indirect association of quality traits owing to a high LD level

In this study, we calculated the LD block using a pairwise comparison of SNP markers separated by 5 Mb. The average LD (*r*^2^ < 0.5) in the parental lines was ∼200 kb (Supplementary Figure 33). The low rate of LD decay in rice means that a GWAS cannot resolve a single gene (Huang et al., 2010). Several significant SNPs were identified in the vicinity of *GW5, GS3, Wx, OsACS6* and *ALK*. Among them, *Wx, OsACS6* and *ALK* influence the ACs of rice grains (Supplementary Figure 34–S36), and *GW5* and *GS3* are closely related to rice grain yield. There are various haplotypes in the V and TC datasets, which may explain why the SNP markers could not achieve statistical significance in those regions (Ye et al., 2018). We ignored other varieties or testcross hybrids containing minor haplotypes. For *Wx* (Fig. 5), Haplotypes A, B and C, which included most varieties in the V dataset, corresponded to haplotypes J, K and L, respectively, in the TC dataset. The varieties having haplotype C had lower ACs (18.50%) and yields (26.15 g) than those having haplotype A or B, while haplotype L accessions had low ACs (18.95%) but also high yields (41.25 g) (Fig. 5a and b). Thus, haplotype L was determined to be optimum for *Wx* in the TC dataset.

**Figure 5.**
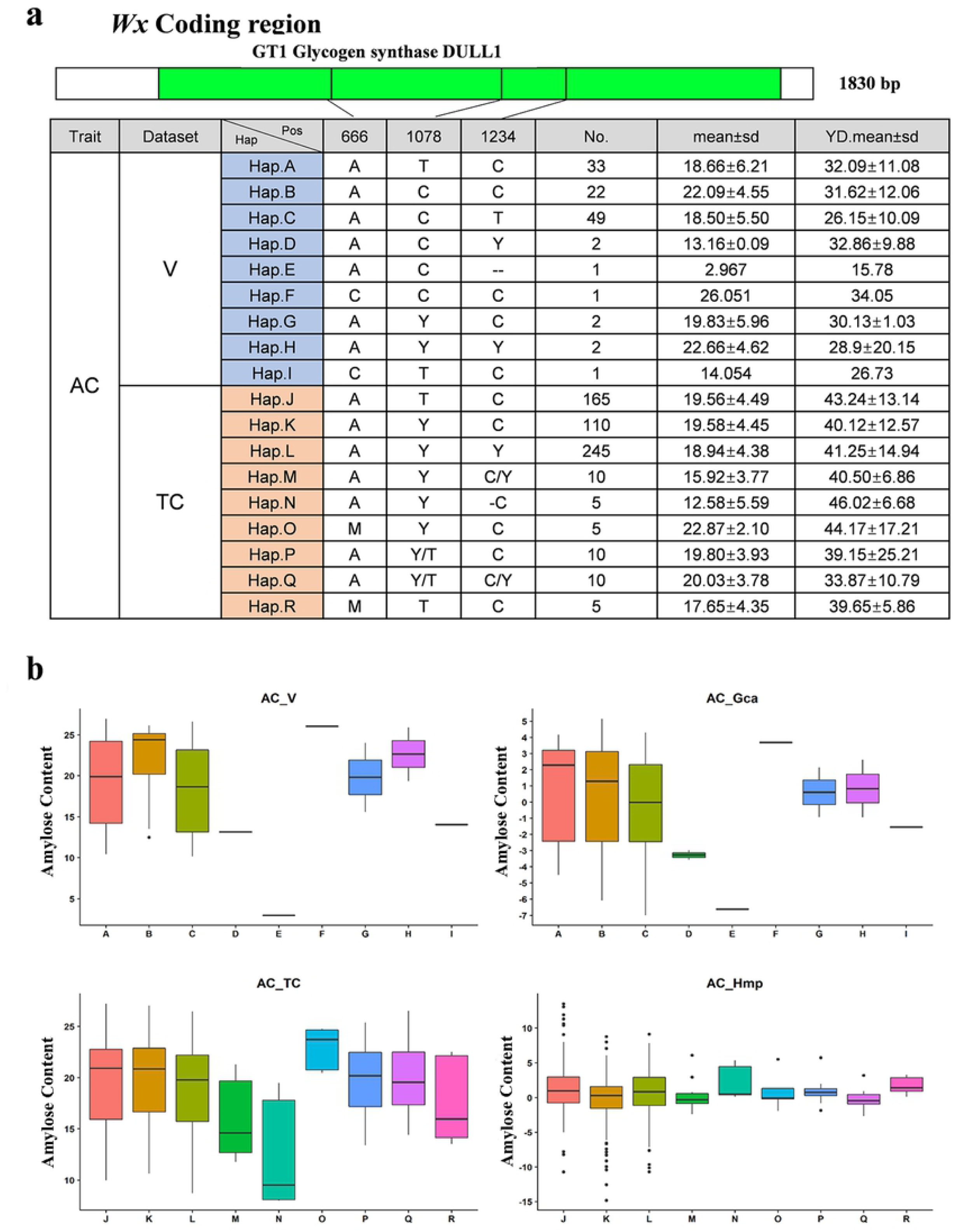
A high degree of polymorphism in the *Wx* coding sequence, and its haplotypes in V and TC datasets. (a) Haplotype analysis of *Wx*. (b) Boxplots for grain width based on the haplotypes of *Wx*.

A similar phenomenon was observed in the coding region of genes *OsACS6* and *ALK*. For *OsACS6*, haplotype M accessions were the optimum (with low ACs and high yields) in the TC dataset, which corresponded to the haplotype B in the V dataset (Supplementary Figure 37). It should be noted that the varieties having haplotype B in the V dataset had low ACs (18.90%) and relatively low yields (23.42 g), but corresponded to haplotype M of the testcross hybrids in the TC dataset, which had low ACs (19.09%) but high yields (37.33 g). Thus, the parental lines with poor performances can still generate elite hybrids owing to heterosis.

For *ALK*, accessions of haplotype DD were optimum in the TC population, corresponding to haplotype D in the V dataset (Supplementary Figure 38). The accessions simultaneously having three superior haplotypes (haplotypes L, M and D for *Wx, OsACS6* and *ALK*, respectively) outperformed (with 18.29% AC and 39.34 g yield) other accessions having only one elite haplotype in the TC dataset.

For *GW5*, accessions of haplotype H were optimum in the TC population, corresponding to haplotype H in the V dataset (Supplementary Figure 39). For *GS3*, accessions having haplotype J were optimum in the TC population, corresponding to haplotype C in the V dataset (Supplementary Figure 40).

### Predicting untested crosses using BayesB

To investigate the quality trait performance of potential hybrids among the 113 inbred lines in our study, we conducted genomic selection predictions using the BayesB method. All the potential hybrids’ predicted values are shown in Supplementary Tables 5–7. The average predicted AC and yield value of the 224 hybrids with accumulating favorable haplotypes of genes (*Wx, OsACS6* and *ALK*) were 17.3% and 42.52 g, which has similar with results from testcross hybrids in this study. When combined them with favorable haplotypes of genes (*GW5* and *GS3*), 18 potential hybrids showed average predicted AC and yield values of 17.06% and 43.16 g. Thus, accumulating favorable haplotypes of important genes related to rice quality and yield traits may help us produce desirable rice hybrids.

## Discussion

In this study, the GWAS method was used to analyze the genetic basis of 12 rice grain quality traits in 113 parental lines and their 565 testcross hybrids. A total of 56 significant SNPs were associated with 9 of the 12 traits in the V, Gca, TC and Hmp datasets. Among them, three significant SNPs were located within or near the cloned quality genes, while the potential roles of the other SNPs in affecting rice grain quality were newly discovered. They provide new information for rice quality trait breeding. A few of these associated loci were detected simultaneously in multiple datasets of the same trait, but most of the significant loci in the different datasets were unique. This indicated that the quality traits’ general combining ability and heterosis are quantitative traits, controlled by multiple genes; however, the phenotypic performance of inbred breeding lines and their general combining ability, as well as phenotypic performance of hybrids and their heterosis, have different genetic bases for rice grain quality. These findings are consisted with the QTL mapping and GWAS analyses of rice agronomic traits presented in previous research using the NCII (north Carolina design II) and TTC (triple test crosses) designs (Li et al. 2008, Qu et al., 2012, Chen et al., 2019).

The dominance degrees of the top 100 associated loci suggested that the phenotypes of quality traits in inbred lines were mainly controlled by A, while the dominance degree distribution trends of grain quality traits of parental lines and testcross hybrids’ phenotypic performances varied in traits, but were mainly controlled by A and PD on the whole. More PD loci existed than other types of loci for GL, GW, GLWR, CD and BRR, while more A loci existed than other type of loci for PGWC, AC, ASV, MRR and HRR. The heteroses of GW, CD, TD and AC (measured by the mid-parental values in this study), GL, PC and BRR, and of GLWR, ASV, MRR and HRR are mainly controlled by OD, PD and A, respectively, which is different from the genetic basis of hybrid rice’ agronomic traits studied by Huang et al (2015). They found fewer OD loci than those with incomplete dominance, which implies that dominance is an important contributor to the heterosis of rice grain quality in hybrid breeding. The average mean values of quality traits increased along with the number of superior alleles of significant associated loci in the four datasets. Similarly, Huang et al. (2015) studied the superior alleles of top 100 GWAS loci (order them in descending of −log(*P* value)) associated with agronomic traits of hybrid progenies and parents. They suggested that the heterosis of hybrid progeny might be superior to that of inbred rice varieties for superior allele accumulation. Our study shows that in terms of quality traits, the phenotypic performance and Gca of inbred rice varieties (in this study, repented as male parental lines), as well as the phenotypic performance and the heterosis of hybrid progeny, may be the result of the accumulation of rare superior alleles. Chen et al. (2019) also found that high Gca parental lines or high Sca (special combining ability) combinations are caused by pyramiding favorable alleles.

Although significant rice grain quality trait-associated loci were detected, pyramiding associated loci of superior alleles does not guarantee improved rice quality. Because of LD, the genes may only be indirectly associated with targeted traits. However, the GWAS results need further analyses, such as transcript and genetic transformational analyses. In this study, the combinations of favorable haplotypes of three indirectly associated genes of AC (*Wx, ALK* and *ACS6*), showed the desired ACs in the male parental lines (18.62%), and surprisingly, the lower AC (15.97%) and higher yield were found in testcross hybrids with the favorable haplotype. Although varieties with combinations of superior alleles of the four significant SNPs associated with AC in the V dataset had low ACs (15.8%), their corresponding lines in the TC dataset had increased ACs, at 17.92% on average. These procedures were also conducted for the traits GL and GW (Fig. 4), and similar trends were observed. Generally, the parental lines with favorable combinations of gene haplotypes (or a favorable haplotype of an individual gene) controlling grain traits crossed with male sterile lines generated hybrids with similar grain quality performances as their male parent but higher yield performances. Thus, hybrids with an improved end-use quality may result from pyramiding favorable gene haplotypes that influence rice quality traits.

Genomic selection predictions of AC, GL, GW and yield for all 6,328 potential hybrids between the 113 inbred lines were conducted. Among the potential hybrids, selecting those with favorable haplotypes of *Wx, ALK, ACS6, GW5* and *GS3* lead to elite ACs (17.06%) and high yields (43.16 g) on average. We are planning to generate these 18 potential hybrids in a future trial experiment. We hope our research may facilitate dissecting the genetic bases of grain quality traits of rice hybrids and help breeders select and produce ideal hybrids.

## Experimental procedures

### Materials and field planting

In accordance with the NCII genetic mating design, 115 *indica* rice accessions, as male parents, were crossed with five male sterile lines, as female parents, to produce testcross hybrids. In 2013, male parental lines (Supplementary Table 1) and testcross hybrids were planted in Wuhan, China. The field trials were designed as randomized blocks, repeated twice, with 20 plants per field planted at a density of 5 × 8 inches. Field management, including irrigation, fertilizer application and pest control, followed normal agricultural practices. The grains were harvested when fully ripe. In total, 113 parental lines (without samples V2 and V3 in Supplementary Table1) and 565 (113 × 5) testcross hybrids were obtained for phenotyping.

### Phenotyping

After the materials matured, three plants with uniform growth were selected from the middle eight plants, dried, and stored at room temperature for 3 months. They were then sent to the Agricultural College of Hunan Agricultural University for rice grain quality evaluation. In accordance with the method of Lou et al. (2009), 12 rice quality traits including the grain length (GL, mm), grain width (GW, mm), GL/GW ratio (GLWR), chalkiness degree (CD, %), percentage of grain with chalkiness (PGWC, %), transparency (transparent degree, TD), amylose content (AC %), alkali spreading value (ASV), PC (total protein content %), brown rice ratio (BRR, %), milled rice ratio(MRR, %) and whole milled rice rate (head rice ratio, HRR, %) of male parental lines and testcross hybrids were determined. The 12 quality traits of female parental lines were measured with those of the five male sterile lines’ seeds. The yield production per plant of male parental lines and testcross hybrids were also evaluated.

### Resequencing, genotyping and imputation

For each parental line, the genomic DNA was prepared from a single plant for sequencing. A sequencing library was established in accordance with the Illumina protocol. The genomes of 118 parental lines were sequenced on an Illumina HiSeq2500 platform with 11× genome coverage on average. ‘Nipponbare’ was used as the reference genome, and BWA software (Li and Durbin, 2009) was used for all paired-end read mapping. Then SNP calling and quality control were conducted by deleting SNPs with minor allele frequency > 5% and missing rate > 20%. In total, 1,894,012 SNPs were obtained and analyzed by NPUTE (ver 4.0, Roberts et al., 2007) with no missing genotype. Testcross hybrid genotypes were inferred using parental SNP genotypic information.

### Phylogenetic and population structures

Neighbor-joining (NJ) trees and principal component analysis (PCA) plots were used to infer the structures of the 118 rice parental lines or 565 testcross hybrids. A pairwise distance matrix derived from the simple matching distance for all the SNP sites was calculated to construct unweighted NJ trees using the software SNPhylo (version 20140116, Lee et al., 2014), and phylogenic trees were draw byiTOL online (http://itol.embl.de/). The program Admixture (ver 1.3) was used to calculate varying levels of *K* (*K* = 1–10; Alexander et al., 2009). The PCA was conducted using GAPIT (version 2.0, Tang et al., 2016) and was separately calculated using SNPs present in parental lines and testcross hybrids. Genome-wide linkage disequilibrium (LD) was estimated using pairwise *r*^2^ values between SNPs, which were calculated using the –r2 -- ld-window99999 --ld-window-r2 0 command in PLINK (version 1.9; Purcell et al., 2007).

### Phenotypic and heterotic statistics and a GWAS analysis

Histograms, boxplots and correlations were constructed using grand means for V or TC. Heterosis was evaluated in testcross hybrids using Hmp, as follows:

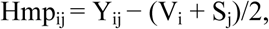

where Y_ij_ represents the phenotypic value of the testcross hybrid derived from the *i*th male parent and *j*th female parent, V_i_ represents the phenotype of the *i*th male parental lines, and *S*_j_ represents the *j*th sterile line phenotypic value. The Gca was measured using the Gca effect (*g*_*i*_), as follows:

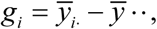

where *g*_*i*_ represents the *i*th maternal parent’s Gca effect, 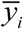. represents the mean phenotypic value of testcross hybrids’ that derived from the *i*th male parent, and 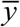 ·· represents the mean phenotypic value of all the testcross hybrids. A GWAS analysis was separately conducted on each trait in the four datasets V, Gca, TC and Hmp using the compressed mixed linear model with the GAPIT software (ver 2.0; Zhang et al., 2010), general and mixed linear models in MVP (https://github.com/XiaoleiLiuBio/rMVP) and FarmCPU (fixed and random model circulating probability unification)(Liu et al., 2016). In addition to the specific parameters of these four algorithms, the PCA was set to 3 for uniformity, and we defined a whole-genome significance cutoff using the Bonferroni test threshold of *P* < 2.64 × 10^−8^ [−log_10_(*P* value) = 7.58, the Bonferroni correction for multiple tests (0.05/n), where n represents the total number of SNPs used in the association analysis].

The TC and Hmp datasets were used for the heterosis analysis. The degree of dominance ‘*d*/*a*’ was calculated using the peak SNPs of the heterosis-associated loci, where ‘*d*’ and ‘*a*’ represent the dominant and the additive effects, respectively. For each significant SNP detected in the TC and Hmp datasets, we denoted the allelic effect of the SNP estimated in the Hmp dataset as the dominant effect and that in the V dataset as the additive effect. This was similar to the method in our previous study (Li et al., 2008). The size of |*d/a*| is the dominance degree, with |*d/a*| > 1.2 indicating overdominance (OD), 0.8 < |*d/a*| ≤ 1.2 indicating dominance (D), 0.2 < |*d/a*| ≤ 0.8 indicating partial dominance (PD) and |*d/a*| ≤ 0.2 indicating additive (A). A GO (GeneOntology) analysis was conducted using candidate genes.

### Superior alleles of the significant associated loci analysis

The effects of heterozygous and homozygous genotypes were calculated for each peak SNP of the associated locus, and the average phenotypic measurements of the heterozygous and homozygous genotypes were calculated. For each significant associated locus, the genotype of the SNP with the preferred quality performance (such as lower AC and greater GL) was set as the superior gene allele (Huang et al., 2015). The numbers of superior alleles of each accession in each dataset were counted.

### Sequence analysis

Because all the parental lines were re-sequenced using the *japonica* cultivar Nipponbare as the references, sequences of some known genes associated with quality traits, such as *OsACS6, Wx, ALK, GW5* and *GS3*, in male parental lines or testcross hybrids were obtained and aligned using the ClustalW method (Thompson et al., 2002).

### Predicting ACs and yields of untested cross hybrids using BayesB method of genomic selection

Following Yang et al.’s method (2019), using the BayesB method, a twenty times fivefold cross-validation was conducted to evaluate the predictability (the correlation coefficient between the observed and predicted phenotypic values) of the rice hybrid AC and yield performances of 565 testcross hybrids using all the SNPs. According to the parameters estimated from the training sample (565 testcross hybrids), AC and yields of all the potential hybrids among the 113 male parental lines were predicted.

## Authorship

LL, XZ, XZ performed data analyses and drafted the manuscript. KX, SS, JS, ZL, WT and ZT assisted in data analysis and discussion. YD, and XZ developed the populations. KX and CW assisted in field data collection. LL and ZY designed and finalized the manuscript, and revised and submitted manuscript. ZH conceived the project, planned, secured extramural funds.

## Acknowledgements

This work was supported by funding from the National key technology research and development program (2016YFD0100101),Open Research Fund of State Key Laboratory of Hybrid Rice (Hunan Hybrid Rice Research Center (2019KF05); Wuhan University (KF201912)), Research Foundation of Education Bureau of Hunan Province (19A244), Double first-class construction project of Hunan Agricultural University (SYL2019028), Hunan Science and Technology Plan Program (2019RS1055), Hubei Provincial cooperative Innovation Center(Hubei Science and Education letter (2016) No.3.

We thank Prof. Sibin Yu and Tongmin Mou from Huazhong Agricultural University, Prof. Shuangcheng Li from Sichuan Agricultural University, Prof. Shuzhu Tang from Yangzhou University, Prof. Guanghua He from Southwest University, Prof. Huaxiong Qi from Hubei Academy of Agricultural Sciences for providing material to us.

## Supplementary Figures

**Supplementary Figure 1.** Phenotype distributions about general combining ability (Gca) of twelve quality traits in parent lines.

**Supplementary Figure 2.** Phenotype distributions about mid-parent heterosis (Hmp) of twelve quality traits in testcross lines.

**Supplementary Figure 3.** Phenotypic correlations among different traits. Correlations were measured in Pearson correlation coefficients. The left bottom panel was phenotype correlations in the TC dataset, and the right top panel was phenotype correlations in the Hmp dataset. Asterisks indicate significant correlations using a two tailed t-test (\**P*<0.05, \*\**P*<0.01 and \*\*\**P*<0.001).

**Supplementary Figure 4.** Genetic architecture of 120 parent lines. (a) Neighbor-joining tree of 115 male parents and 5 female parents used to construct the NCII population. (b) Genetic structure of parental lines analyzed using the program admixture.

**Supplementary Figure 5.** Genetic architecture of testcross lines. (a) Neighbor-joining tree of 565 testcross lines. (b) Genetic structure of testcross lines analyzed using the program admixture.

**Supplementary Figure 6.** Plots of first two component in 120 parental lines.

**Supplementary Figure 7.** Plots of first four component in 565 testcross lines. (a) Plot of first and second components in testcross lines; (b) Plot of third and fourth component in testcross lines; (c) Plot of first three components in testcross lines.

**Supplementary Figure 8.** Manhattan plots of GL. (a) Manhattan plots of GL_V using four different algorithms; (b) Manhattan plots of GL_Gca using four different algorithms; (c) Manhattan plots of GL_TC using four different algorithms; (d) Manhattan plots of GL_Hmp using four different algorithms.

**Supplementary Figure 9.** QQ-plots of GL. (a) QQ-plots of GL_V using four different algorithms; (b) QQ-plots of GL_Gca using four different algorithms; (c) QQ-plots of GL_TC using four different algorithms; (d) QQ-plots of GL_Hmp using four different algorithms.

**Supplementary Figure 10.** Manhattan plots of GW. (a) Manhattan plots of GW_V using four different algorithms; (b) Manhattan plots of GW_Gca using four different algorithms; (c) Manhattan plots of GW_TC using four different algorithms; (d) Manhattan plots of GW_Hmp using four different algorithms.

**Supplementary Figure 11.** QQ-plots of GW. (a) QQ-plots of GW_V using four different algorithms; (b) QQ-plots of GW_Gca using four different algorithms; (c) QQ-plots of GW_TC using four different algorithms; (d) QQ-plots of GW_Hmp using four different algorithms.

**Supplementary Figure 12.** Manhattan plots of GLWR. (a) Manhattan plots of GLWR_V using four different algorithms; (b) Manhattan plots of GLWR_Gca using four different algorithms; (c) Manhattan plots of GLWR_TC using four different algorithms; (d) Manhattan plots of GLWR_Hmp using four different algorithms.

**Supplementary Figure 13.** QQ-plots of GLWR. (a) QQ-plots of GLWR_V using four different algorithms; (b) QQ-plots of GLWR_Gca using four different algorithms; (c) QQ-plots of GLWR_TC using four different algorithms; (d) QQ-plots of GLWR_Hmp using four different algorithms.

**Supplementary Figure 14.** Manhattan plots of CD. (a) Manhattan plots of CD_V using four different algorithms; (b) Manhattan plots of CD_Gca using four different algorithms; (c) Manhattan plots of CD_TC using four different algorithms; (d) Manhattan plots of CD_Hmp using four different algorithms.

**Supplementary Figure 15.** QQ-plots of CD. (a) QQ-plots of CD_V using four different algorithms; (b) QQ-plots of CD_Gca using four different algorithms; (c) QQ-plots of CD_TC using four different algorithms; (d) QQ-plots of CD_Hmp using four different algorithms.

**Supplementary Figure 16.** Manhattan plots of PGWC. (a) Manhattan plots of PGWC_V using four different algorithms; (b) Manhattan plots of PGWC_Gca using four different algorithms; (c) Manhattan plots of PGWC_TC using four different algorithms; (d) Manhattan plots of PGWC_Hmp using four different algorithms.

**Supplementary Figure 17.** QQ-plots of PGWC. (a) QQ-plots of PGWC_V using four different algorithms; (b) QQ-plots of PGWC_Gca using four different algorithms; (c) QQ-plots of PGWC_TC using four different algorithms; (d) QQ-plots of PGWC_Hmp using four different algorithms.

**Supplementary Figure 18.** Manhattan plots of TD. (a) Manhattan plots of TD_V using four different algorithms; (b) Manhattan plots of TD_Gca using four different algorithms; (c) Manhattan plots of TD_TC using four different algorithms; (d) Manhattan plots of TD_Hmp using four different algorithms.

**Supplementary Figure 19.** QQ-plots of TD. (a) QQ-plots of TD_V using four different algorithms; (b) QQ-plots of TD_Gca using four different algorithms; (c) QQ-plots of TD_TC using four different algorithms; (d) QQ-plots of TD_Hmp using four different algorithms.

**Supplementary Figure 20.** Manhattan plots of AC. (a) Manhattan plots of AC_V using four different algorithms; (b) Manhattan plots of AC_Gca using four different algorithms; (c) Manhattan plots of AC_TC using four different algorithms; (d) Manhattan plots of AC_Hmp using four different algorithms.

**Supplementary Figure 21.** QQ-plots of AC. (a) QQ-plots of AC_V using four different algorithms; (b) QQ-plots of AC_Gca using four different algorithms; (c) QQ-plots of AC_TC using four different algorithms; (d) QQ-plots of AC_Hmp using four different algorithms.

**Supplementary Figure 22.** Manhattan plots of ASV. (a) Manhattan plots of ASV_V using four different algorithms; (b) Manhattan plots of ASV_Gca using four different algorithms; (c) Manhattan plots of ASV_TC using four different algorithms; (d) Manhattan plots of ASV_Hmp using four different algorithms.

**Supplementary Figure 23.** QQ-plots of ASV. (a) QQ-plots of ASV_V using four different algorithms; (b) QQ-plots of ASV_Gca using four different algorithms; (c) QQ-plots of ASV_TC using four different algorithms; (d) QQ-plots of ASV_Hmp using four different algorithms.

**Supplementary Figure 24.** Manhattan plots of PC. (a) Manhattan plots of PC_V using four different algorithms; (b) Manhattan plots of PC_Gca using four different algorithms; (c) Manhattan plots of PC_TC using four different algorithms; (d) Manhattan plots of PC_Hmp using four different algorithms.

**Supplementary Figure 25.** QQ-plots of PC. (a) QQ-plots of PC_V using four different algorithms; (b) QQ-plots of PC_Gca using four different algorithms; (c) QQ-plots of PC_TC using four different algorithms; (d) QQ-plots of PC_Hmp using four different algorithms.

**Supplementary Figure 26.** Manhattan plots of BRR. (a) Manhattan plots of BRR_V using four different algorithms; (b) Manhattan plots of BRR_Gca using four different algorithms; (c) Manhattan plots of BRR_TC using four different algorithms; (d) Manhattan plots of BRR_Hmp using four different algorithms.

**Supplementary Figure 27.** QQ-plots of BRR. (a) QQ-plots of BRR_V using four different algorithms; (b) QQ-plots of BRR_Gca using four different algorithms; (c) QQ-plots of BRR_TC using four different algorithms; (d) QQ-plots of BRR_Hmp using four different algorithms.

**Supplementary Figure 28.** Manhattan plots of MRR. (a) Manhattan plots of MRR_V using four different algorithms; (b) Manhattan plots of MRR_Gca using four different algorithms; (c) Manhattan plots of MRR_TC using four different algorithms; (d) Manhattan plots of MRR_Hmp using four different algorithms.

**Supplementary Figure 29.** QQ-plots of MRR. (a) QQ-plots of MRR_V using four different algorithms; (b) QQ-plots of MRR_Gca using four different algorithms; (c) QQ-plots of MRR_TC using four different algorithms; (d) QQ-plots of MRR_Hmp using four different algorithms.

**Supplementary Figure 30.** Manhattan plots of HRR. (a) Manhattan plots of HRR_V using four different algorithms; (b) Manhattan plots of HRR_Gca using four different algorithms; (c) Manhattan plots of HRR_TC using four different algorithms; (d) Manhattan plots of HRR_Hmp using four different algorithms.

**Supplementary Figure 31.** QQ-plots of HRR. (a) QQ-plots of HRR_V using four different algorithms; (b) QQ-plots of HRR_Gca using four different algorithms; (c) QQ-plots of HRR_TC using four different algorithms; (d) QQ-plots of HRR_Hmp using four different algorithms.

**Supplementary Figure 32.** The dominance degree of 12 quality traits. (a) The dominance of V in each trait; (b) The dominance of Gca in each trait; (c) The dominance of TC in each trait; (d) The dominance of Hmp in each trait.

**Supplementary Figure 33.** LD decay in 120 parental lines.

**Supplementary Figure 34.** Analyses of the peak for amylose content (near gene *Wx*) associated haplotypes at chromosome 6 in TC. Local Manhattan plot (top) and LD heatmap (bottom) surrounding the peak on chromosome 6. The red dash line indicated the positions of of *Wx*.

**Supplementary Figure 35.** Analyses of the peak for amylose content (near gene *OsACS6s*) associated haplotypes at chromosome 6 in V (a) and TC dataset (b). Local Manhattan plot (top) and LD heatmap (bottom) surrounding the peak on chromosome 6. The red dash line indicated the positions of of *OsACS6s*.

**Supplementary Figure 36.** Analyses of the peak for amylose content (near gene *ALK*) associated haplotypes at chromosome 6 in V (a) and TC dataset (b). Local Manhattan plot (top) and LD heatmap (bottom) surrounding the peak on chromosome 6. The red dash line indicated the positions of of *ALK*.

**Supplementary Figure 37.** A high degree of polymorphism in the OsACS6 coding sequence, and its haplotypes in V and TC datasets. (a) Haplotype analysis of OsACS6. (b) Boxplots for grain width based on the haplotypes of OsACS6.

**Supplementary Figure 38.** A high degree of polymorphism in the ALK coding sequence, and its haplotypes in V and TC datasets. (a) Haplotype analysis of ALK. (b) Boxplots for grain width based on the haplotypes of ALK.

**Supplementary Figure 39.** A high degree of polymorphism in the GW5 coding sequence, and its haplotypes in V and TC datasets. (a) Haplotype analysis of GW5. (b) Boxplots for grain width based on the haplotypes of GW5.

**Supplementary Figure 40.** A high degree of polymorphism in the GS3 coding sequence, and its haplotypes in V and TC datasets. (a) Haplotype analysis of GS3. (b) Boxplots for grain width based on the haplotypes of GS3. (c) Unrooted phylogenetic tree of GS3.

## Supplementary Tables

**Supplementary Table 1.** The information of 115 varieties and 5 sterile lines used in NCII design.

**Supplementary Table 2.** Selected model of GWAS for 12 rice quality traits.

**Supplementary Table 3.** The information of SNP pleiotropism.

**Supplementary Table 4.** The average value of each trait in different number of Superior alleles.

**Supplementary Table 5.** Untested cross AC performance and their SNP genotype of associated loci or haplotype of gene.

**Supplementary Table 6.** Untested cross GL performance and their SNP genotype of associated loci or haplotype of gene.

**Supplementary Table 7.** Untested cross GW performance and their SNP genotype of associated loci or haplotype of gene.

**Supplementary Table 8.** Phenotype data of parental lines.

**Supplementary Table 9.** Phenotype data of testcross lines.

